# Development of genomic markers associated to production traits in lumpfish (*Cyclopterus lumpus*)

**DOI:** 10.1101/2023.05.10.540148

**Authors:** Alejandro P. Gutierrez, Sarah-Louise Counter Selly, John B. Taggart, Panagiotis Kokkinias, Thomas Cavrois-Rogacki, Eduardo Jimenez Fernandez, Herve Migaud, Ingrid Lein, Andrew Davie, Michaël Bekaert

**Affiliations:** Institute of Aquaculture, University of Stirling, Stirling FK9 4LA, Scotland, UK; Otter Ferry SeaFish Ltd., Otter Ferry, Tighnabruaich, PA21 2DH, Scotland, UK; Nofima AS, Sjølsengvegen 22, 6600 Sunndalsøra, Norway

**Keywords:** Lumpfish, QTL, GWAS, Aquaculture

## Abstract

Cleaner fish species have gained great importance in the control of sea lice, among them, lumpfish (*Cyclopterus lumpus*) has become one of the most popular. Lumpfish life cycle has been closed, and hatchery reproduction is now possible, however, current production is reliant on wild caught broodstock to meet the increasing demand. Selective breeding practices are called to play an important role in the successful breeding of most aquaculture species, including lumpfish.

In this study we analysed a lumpfish population for the identification of genomic markers linked to production traits. Sequencing of RAD libraries allowed us to identify, 7,193 informative markers within the sampled individuals. Genome wide association analysis for sex, weight, condition factor and standard length was performed. One single major QTL region was identified for sex determination, while nine QTL regions were detected for weight, and three QTL regions for standard length.

A total of 177 SNP markers of interest (from QTL regions) and 399 top F_*st*_ SNP markers were combined in a low-density panel, useful to obtain relevant genetic information from lumpfish populations. Moreover, a robust combined subset of 29 SNP markers (10 associated to sex, 14 to weight and 18 to standard length) provided over 90% accuracy in predicting the animal’s phenotype. Overall, our findings provide significant insights into the genetic control of important traits in lumpfish and deliver important genomic resources that will facilitate the establishment of selective breeding programs in lumpfish.

## INTRODUCTION

Sea lice infestation remains the most pressing issue affecting salmon aquaculture worldwide. Losses linked to sea lice were estimated at C700 million worldwide in 2015 and continues to increases (Brooker et al., 2018). These losses not only result from reduced production due to sea lice-associated mortalities, decreased fish growth, and reduced flesh quality, but also from the cost of treatment against sea lice. This often involves the use of parasiticide chemicals or mechanical treatments of limited effectiveness or carry other health risks (Costello, 2009).

To address this issue, biological control of sea lice infection in Atlantic salmon cages has become an important alternative to tackle the one of the most important diseases affecting salmon aquaculture. This strategy has gained increasing popularity mainly due to its effectiveness and environmental safety (Costello, 2006). In Norway, for example, about 0.7 million cleaner fish were deployed in salmon cages in 2006 which drastically increased to 43 million cleaner fish deployed in 2019, while showing a slight decrease in the last couple of years (Norwegian Directorate of Fisheries, 2022). Estimated figures indicate that approximately half of the cleaner fish used in the Atlantic salmon industry are lumpfish (*Cyclopterus lumpus*), while the remainder include various species of wrasse (Bolton-Warberg, 2018; Overton et al., 2020) such as ballan wrasse (*Labrus bergylta*), corkwing wrasse (*Symphodus melops*) and Goldsinny wrasse (*Ctenolabrus rupestris*). However, current cleaner fish production relies heavily on wild-caught broodstock, and production from farmed broodstock remains minimal (Powell et al., 2018; Bolton-Warberg, 2018).

In recent years, hatchery production technologies for cleaner fish, particularly lumpfish and ballan wrasse, have been under intense development. Among these species, lumpfish hatchery production has proven to be more straightforward, with relatively high and stable survival rates (Brooker et al., 2018) being achieved, making it a promising candidate for extensive use as a cleaner fish (Imsland et al., 2018). Lumpfish is a sub-Arctic species commonly found along the Icelandic, Norwegian, and British coastlines, as well as the East coast of North America (Davenport, 1985). Crucially, the lumpfish life cycle has been closed, and hatchery reproduction is now possible. However, there are only a limited number of hatcheries producing lumpfish in Europe, and breeding programs are notably lacking (Brooker et al., 2018). Selective breeding is an effective strategy for improving the production of aquaculture species, by enhancing economically important traits (Regan et al., 2021). Mimicking natural conditions, lumpfish hatchery production takes only around seven months until fish are ready for deployment, much shorter than the approximatively 1.5 years currently needed for ballan wrasse deployment, making the production cycle considerably more economically viable (Brooker et al., 2018). Contrary to most aquaculture species, lumpfish grow very faster than would be preferred, leading to problems associated with delousing behaviour (Imsland et al., 2016). There is a marked decline in delousing activity in lumpfish upon reaching a large body size (over 300 g in 6-10 months), likely due to their slow movement, feeding off salmon pellet, and increasing aggressive (and territorial) behaviour, mainly triggered by the onset of sexual maturation (Imsland et al., 2016). Therefore, the establishment of breeding programmes that allow the production of stocks with a more favourable growth rate and other desirable traits would greatly benefit lumpfish production.

Recent developments in genomic technologies have transformed selective breeding programs for aquaculture species. This has mostly been driven by continuous advances in sequencing technologies that enable high-throughput discovery and screening of genetic markers, in particular single nucleotide polymorphisms (SNPs), which are highly abundant and widely distributed through the genome (Houston et al., 2020). Screening of thousands of SNPs via genotyping-by-sequencing (GBS) techniques or by SNP arrays has become common practise in devising and managing selective breeding schemes for many commercial aquaculture species, including emerging ones (Houston et al., 2020; Robledo et al., 2018). In addition, reference genome assemblies have been developed for numerous aquaculture species (Yue and Wang, 2017), providing a keystone for advanced genetic studies of their biology and potential improvement. Accordingly, the recent release of a reference genome for lumpfish serves as a valuable genomic resource for advancing production of this species (Holborn et al., 2022).

To support the establishment of sustainable selective breeding programmes for lumpfish, this study aimed to develop genomic resources for lumpfish and identify genomic regions associated with growth traits and sex. These efforts will aid in the production of stocks with favourable growth rates and other traits of interest, ultimately contributing to the development of an effective and sustainable solution to the sea lice infestation issue in salmon aquaculture.

## MATERIALS AND METHODS

### Family creation

Wild broodstock were obtained from Skjerneset Fisk at Averøy, Norway. A total of 14 *C. lumpus* independent families (1Q to 14Q) were created, within 3 hours of each other on the same day in October 2018, and reared from fertilisation to final sampling at the NOFIMA Cleaner Fish Unit at Sunndalsora, Norway. All families were reared in discrete incubation units/tanks that were supplied by a common water source to ensure comparable environmental conditions. Hatching in all families initiated within three calendar days of each other, no later than 300-degree days post fertilisation. Larvae were fed following routine commercial practice, first with live feed (Artemia), before weaning to a commercial formulated feed (Otohime, PTAqua, Norway). At 90 days post hatch, when fish reached an average of 0.58 g wet weight, the total number of families being reared was rationalised to four and stock numbers balanced to an average of 3,250 juveniles which were selected at random. The final four families were reared following normal commercial practice until 180 days post hatch, when the final phenotyping sampling was performed.

### Phenotype Measurement

For all four families the same sampling regime was followed. A total of 100 individuals were selected at random, culled by lethal anaesthesia and then for each individual total length (*±*1 mm), standard length (*±*1 mm), weight (*±*0.01 g) and sex (where identifiable) were measured. There-after, a family specific upper and lower size threshold was calculated (bottom 10% of population curve “small” and top 10% of population curve “big”). Thereafter a further 100 “small” and “big” individuals within each family were sampled as shown in Table 1. For all fish body condition was measured using Fulton’s condition factor (K = 100 *×* weight/length3; (Nash et al., 2006)). The subsequent genomic analysis was based on a selection of 50 “big”, 50 “small” as well as 25 random selected individuals from each family. Gender of juveniles within each family was balanced where possible.

**Table 1.** Summary of population statistics. Mean weight (g) and length (mm) distributions as well as maximum and minimum sizes observed, along with threshold sizes (as defined by individual total length) which demarked population specific upper (90%) and lower (10%) size thresholds for selective sampling.

|  | Wet weight (g) | Total Length (mm) | Largest<br>Individual | Smallest<br>Individual | 90% & 10%<br>threshold |
| --- | --- | --- | --- | --- | --- |
| Family 3Q | 9.76 ± 7.7 | 57.59 ± 13.4 | 57.13 g<br>105 mm | 0.81 g<br>30 mm | > 71 mm<br>< 41 mm |
| Family 10Q | 5.44 ± 5.2 | 46.99 ± 12.5 | 51.25 g<br>107 mm | 0.47 g<br>25 mm | > 66 mm<br>< 33 mm |
| Family 11Q | 3.64 ± 5.7 | 39.84 ± 12.5 | 43.82 g<br>97 mm | 0.25 g<br>21 mm | > 57 mm<br>< 29 mm |
| Family 7bQ | 2.79 ± 0.8 | 39.53 ± 8.7 | 20.14 g<br>77 mm | 0.36 g<br>22 mm | > 50 mm<br>< 31 mm |

### DNA Extraction

Fin clips for all parents and offspring were stored in 99% ethanol at 4 °C until DNA extraction. Genomic DNA was extracted using a salt extraction method as described before (Brown et al., 2016). Total nucleic acid content and quality (260 nm/230 nm and 260 nm/280 nm ratios) were determined by spectrometry (Nanodrop; Thermo Scientific, Hemel Hempstead, UK) before measuring double-stranded DNA concentrations using a Qubit dsDNA Broad Range Assay Kit and Qubit Fluorometer (Invitrogen, Paisley, UK).

### Library Preparation and Sequencing

The ddRAD libraries were prepared using an adapted version of an existing protocol (Brown et al., 2016). Briefly, DNA from each sample was digested at 37 °C, for 75 min with restriction enzymes *Pst*I and *Nla*III (New England Biolabs, UK), followed by heat-inactivation at 65 °C, for 25 min. The DNA samples were then individually barcoded through the ligation of specific P1 and P2 adapters, each containing a unique five or seven base nucleotide sequence. After addition of pre-mixed adaptors (*Pst*I:*Nla*III 1:16) and incubation of samples at 22 °C, for 10 min, T4 ligase (2000 ceU/*µ*g DNA), rATP (100 mM) and CutSmart buffer (1*×*) were added and samples incubated for 90 min at 22 °C, followed by heat inactivation (65 °C, 20 min). Libraries were column purified (PCR MinElute, Qiagen, Manchester, UK), size selected by gel electrophoresis (550-650 bp) and amplified by PCR (15 cycles). Sequencing was performed by Novogene (UK) Co. Ltd. (Cambridge, UK) using an Illumina NovaSeq 6000 platform (150-base paired-end reads).

### Marker Assembly and Genotyping

The sequence data from the 536 individuals (Supplementary Table S1) were pre-processed to discard low quality reads (i.e., with an average quality score less than 20). Sequences lacking the restriction site or having ambiguous barcodes were discarded during sample demultiplexing stage. Retained reads were then aligned against the genomic assembly of *C. lumpus* (NCBI Assembly accession GCA 009769545.1) using bowtie2 v2.3.5.1 (Langmead and Salzberg, 2012) and assembled using *gstacks* from Stack v2.60 (Catchen et al., 2011).

All loci that were common to at least two individuals, with no further filtering, were exported from Stacks. Using PLINK v1.9 (Purcell et al., 2007), groups of variants that shared the same coordinates were identified, and only the first marker was retained (*–list-duplicate-vars suppress-first*), to avoid duplications or indistinctions. Moreover, SNPs with unknown position or located in partial chromosomes were excluded from the analysis. For each dataset (parents and offspring), SNPs and individuals were further filtered for quality control in a two steps process, again using PLINK. First, SNP inclusion was confined to those with minor allele frequency ≥ 0.005 and *p*-value of *χ*^2^ test for Hardy-Weinberg equilibrium ≥ 10-6. Then SNPs and animals with a call rate ≥ 0.9 were selected. Quality control was performed on the datasets (parents and offspring) independently. Filtered scores were then combined in one dataset, keeping only shared SNPs.

### Multidimensional Scaling Analysis

R v4.2.0 (R Core Team, 2022) was used to carry out Multidimensional Scaling Analysis on the dataset using the package Bioconductor/SNPRelate v1.30.1 (Zheng et al., 2012) to calculate the Identity-By-State (IBS) proportion for each sample.

### Identification of Trait Associated Markers

Using the recorded phenotypic data (total length, standard length, weight, Condition index and sex) association analyses were performed within the package R/SNPassoc v2.0-11 (González et al., 2007), using the “log-additive” model (except for sex, where ‘co-dominant” model was used) and R/qtl2 v0.28 (Broman et al., 2018) for R v4.2.0 (R Core Team, 2022). We used a *p*-value threshold of 0.001 and a corrected *p*-value for multiple tests of 0.001/*number of tests*. The model used for the analysis was based on Interval Mapping. The algorithm used considers the phenotype to follow a mixture of Bernoulli distributions and uses a form of the EM algorithm for obtaining maximum likelihood estimates (Broman and Sen, 2009). Two-way and multiple quantitative trait locus (QTL) models were also run with this package.

### QTL Strength Model

The effect of all SNP markers for each QTL was analysed using WEKA v3.8 (Witten et al., 2017), which contains a variety of machine-learning algorithms, including “REPTree” (Witten et al., 2017), a fast decision tree learner that builds a decision/regression tree using information gain/variance and reduced-error pruning with backfitting. “REPTree” considers all the markers, then derives for each individual a phenotype prediction (lengths, weight and sex) based on its genotypes for the markers considered. The most predictive SNP markers for each QTL were selected and used to produce a reduced SNP panel with the same prediction power compared to the full set of markers using WEKA v3.8 (Witten et al., 2017). Permutatively, individuals were removed one-by-one from the training set, with the algorithm subsequently assigning their predicted phenotypic values.

### Low Density SNP Panel

To develop an extensive SNP panel able to capture all genomic regions of interest, as well as maximising the estimation of diversity, all SNP markers associated with the phenotypes of interest (lengths, weight, and sex) were selected as well as markers with the highest F_*st*_ values. F_*st*_ were calculated using the function *gl*.*fst*.*pop* from dartR v2.7.2 (Gruber et al., 2018) and based on all available samples/families. After several tests run by LGC Genomics (Teddington, UK), SNP markers that presented successful amplification by SeqSNP were used for the final panel.

### Panel Validation

The usefulness of the SNP panel was validated to confirm the association of the selected markers to the analysed traits. For this purpose, additional members of the four families previously used in the genome-wide association study (GWAS) analyses, as well as parents, were genotyped, selecting the rest of the “big” and “small” individuals from each family (Supplementary Table S2). In total, tissue samples from 477 fish were shipped and genotyped by LGC Genomics (Teddington, UK).

## RESULTS

### Library Sequencing

High throughput sequencing of 536 individuals produced 3,260,920,744 paired-end reads in total. After the removal of low-quality and incomplete reads, 78.9% of the total raw reads were retained (2,571,378,028 PE-reads; Supplementary Table S1). *C. lumpus* genome was used to map the reads and generate ddRAD-tags. A total of 3,048,066 unique loci were detected, with 477,421 shared by at least two samples.

### SNP Identification and Quality Control

From the 477,421 SNPs initially identified between the two groups (36 parents and 500 offspring), the filtering process left 35 parents with 19,227 SNPs passing the threshold, and 499 offspring with 8,186 SNPs, as shown in Table 2. A total of 7,193 common informative markers were identified (covering the remaining 534 individuals) and used in subsequent analyses (Supplementary Data S1 and Supplementary Table S3).

**Table 2.**
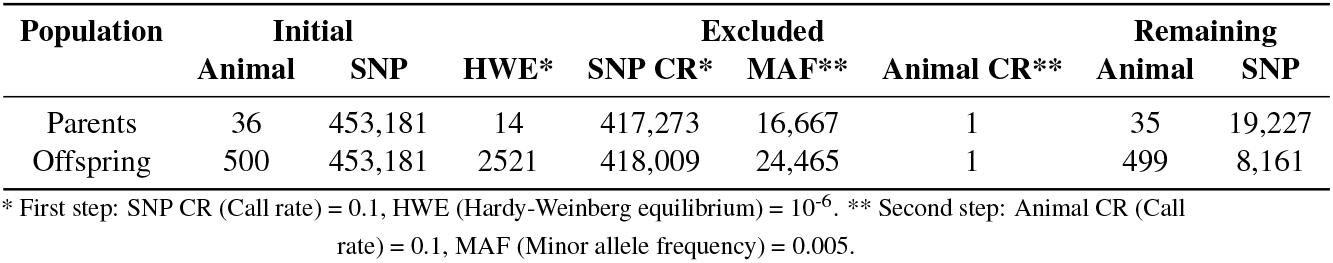
SNP markers filtering steps.

### Sample Structure

A Multidimensional Scaling Analysis (IBS) was utilised to capture the complex structure of the samples and separate the individuals into clusters based on their genetic distance (Jolliffe and Cadima, 2016). This process grouped individuals of same origins together (families), while positioning prior family assignment errors or poor-quality samples as outliers (Figure 1). Five distinct clusters were separated using the first two components (67.3% of cumulative variance). Families and parental/wild generation were clearly clustered. There was one exception; individual 11Q-212 did not behave as expected and did not cluster with any of the families, most likely due to wrong assignment during sampling or handling issues during the transfer into family tanks.

**Figure 1.**
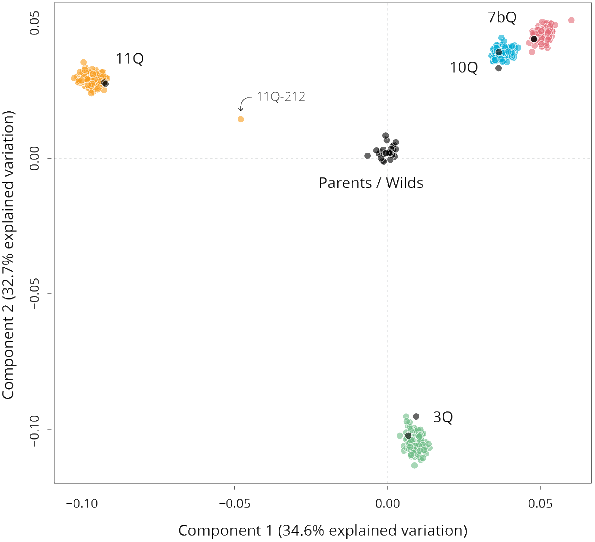
Multidimensional Scaling Analysis of the full dataset. The first and second components explain 34.6%, and 32.7% of the variation found. Based on 7,193 SNP markers.

### Association Analysis

Making use of the 7,193 QC filtered and informative SNP markers, R/SNPassoc and R/qtl2 were used to conduct a QTL/GWAS analysis for both sex and morphometric ratios measurements. Genome wide association was detected for both sex and morphometric measurements after Bonferroni test correction (Supplementary Table S3). One single major QTL (57 SNPs) was identified for sex determination (Figure 2a), whereas a total of nine QTL regions (120 SNPs) were detected for weight (Figure 2b), and three QTL regions (23 SNPs) for standard length (Figure 2c). On the other hand, no significant association was detected when using Fulton’s condition factor as a trait (Figure 2d). All SNP markers associated with standard length were also significantly associated with Weight (Figure 3 and Table 3).

**Table 3.** QTL detected in this study and the genomic regions harbouring them. The peaks and confidence intervals (CI) can be visualised in Figure 2, while the QTLs are reported in Figure 3.

| Traits | QTL | Chr. | CI (bp) | Peak (LOD; bp) | Markers | Mean LOD |
| --- | --- | --- | --- | --- | --- | --- |
| Sex | SexQTL | 13 | 888,322 – 16,609,738 | 29.5; 5,952,081 | 57 | 12.8 |
| Weight | QTL1* | 1 | 3,584,426 – 9,963,809 | 8.3; 3,584,426 | 2 | 8.3 |
|  | QTL2 | 2 | 1,931,736 – 1,931,736 | 8.3; 1,931,736 | 1 | 8.3 |
|  | QTL2 | 3 | 4,805,596 – 26,754,368 | 10.2; 26,754,368 | 12 | 8.1 |
|  | QTL4 | 4 | 11,522,283 – 23,529,190 | 8.3; 19,116,224 | 13 | 7.3 |
|  | QTL5 | 5 | 3,362,532 – 22,429,695 | 9.3; 4,965,412 | 15 | 8.3 |
|  | QTL6* | 6 | 1,992,512 – 5,395,107 | 8.2; 1,992,512 | 2 | 7.9 |
|  | QTL7* | 8 | 20,552,086 – 20,552,086 | 7.1; 20,552,086 | 1 | 7.1 |
|  | QTL8* | 9 | 13,608,747 – 25,957,741 | 8.3; 13,608,747 | 2 | 7.6 |
|  | QTL9 | 10 | 249,201 – 17,391,982 | 8.3; 249,201 | 4 | 7.7 |
|  | QTL10 | 11 | 13,911,073 – 22,899,415 | 8.3; 1,6341,550 | 5 | 7.4 |
|  | QTL11 | 12 | 4,710,808 – 18,097,392 | 7.9; 18,097,392 | 9 | 7.6 |
|  | QTL12* | 13 | 8,434,979 – 16,572,657 | 8.0; 16,572,657 | 3 | 7.7 |
|  | QTL13 | 14 | 260,826 – 23,505,589 | 9.4; 23,505,589 | 15 | 8.1 |
|  | QTL14* | 15 | 6,396,440 – 6,396,440 | 7.2; 6,396,440 | 1 | 7.2 |
|  | QTL15* | 17 | 11,464,082 – 13,337,808 | 8.3; 11,464,082 | 2 | 8.1 |
|  | QTL16* | 18 | 8,262,460 – 13,642,938 | 8.3; 8,262,460 | 2 | 8.3 |
|  | QTL17 | 19 | 470,829 – 16,571,612 | 13.2; 1,188,533 | 17 | 10.4 |
|  | QTL18* | 21 | 14,317,822 – 24,347,228 | 7.9; 24,347,228 | 3 | 7.6 |
|  | QTL19 | 22 | 5,460,636 – 21,864,352 | 9.4; 16,287,751 | 10 | 8.6 |
|  | QTL20* | 25 | 11,069,039 – 11,069,039 | 7.5; 11,069,039 | 1 | 7.5 |
| Std length | QTL1 | 5 | 4,965,412 – 13,898,772 | 7.6; 4,965,412 | 6 | 7.3 |
|  | QTL2* | 14 | 23,505,589 – 23,505,589 | 7.1; 23,505,589 | 1 | 7.1 |
|  | QTL3 | 19 | 470,829 – 1,3758,119 | 11.7; 1,188,533 | 12 | 10.2 |
|  | QTL4 | 22 | 1,6287,751 – 21,864,352 | 7.4; 19,726,257 | 4 | 7.1 |

\* Small peaks not reported in Figure 3.

**Figure 2.**
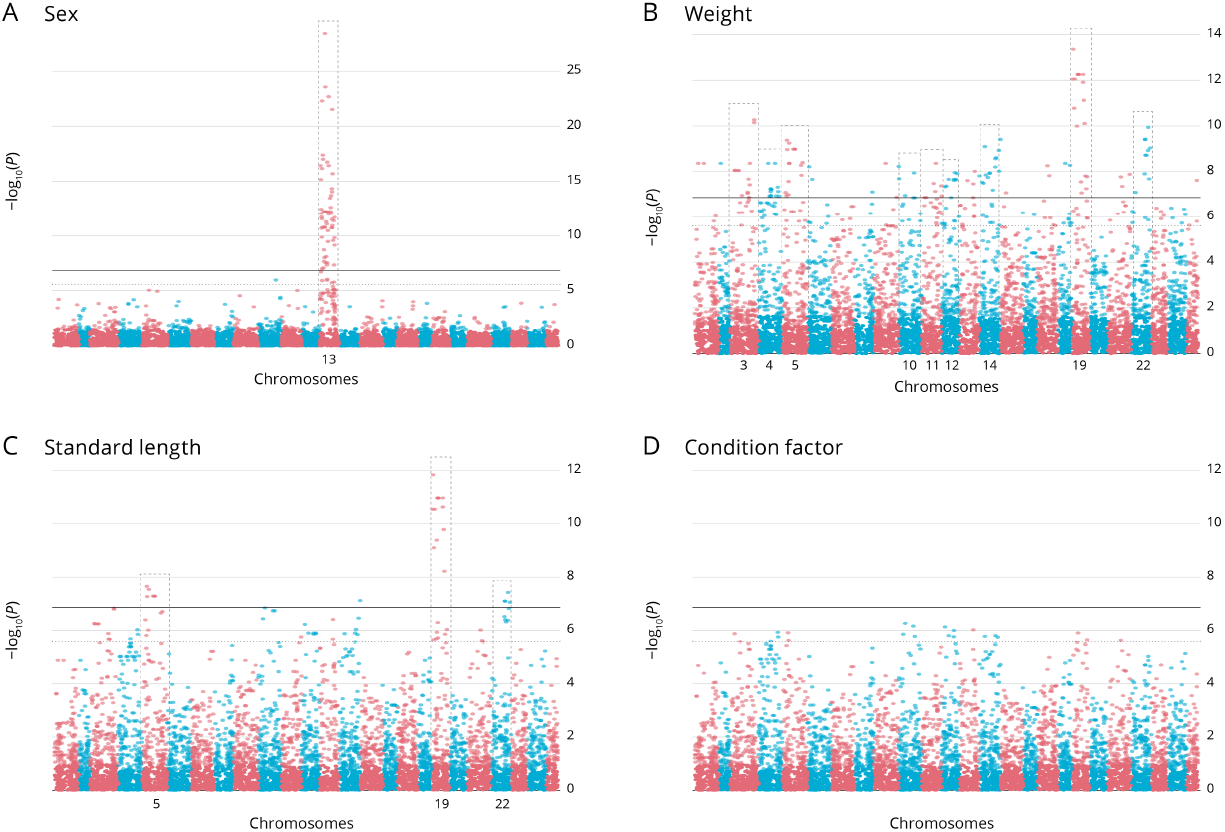
Markers associated with phenotypes. (**a**) Manhattan plot of the association for phenotypic sex. (**b**) Manhattan plot of the association with the fish weight. (**c**) Manhattan plot of the association with the fish Standard length. (**d**) Manhattan plot of the association with the fish Condition factor. The -log_10_(*p*-value) values for association of directly genotyped SNPs are plotted as a function of position of the genetic map. Each chromosome has been represented with a different colour.

**Figure 3.**
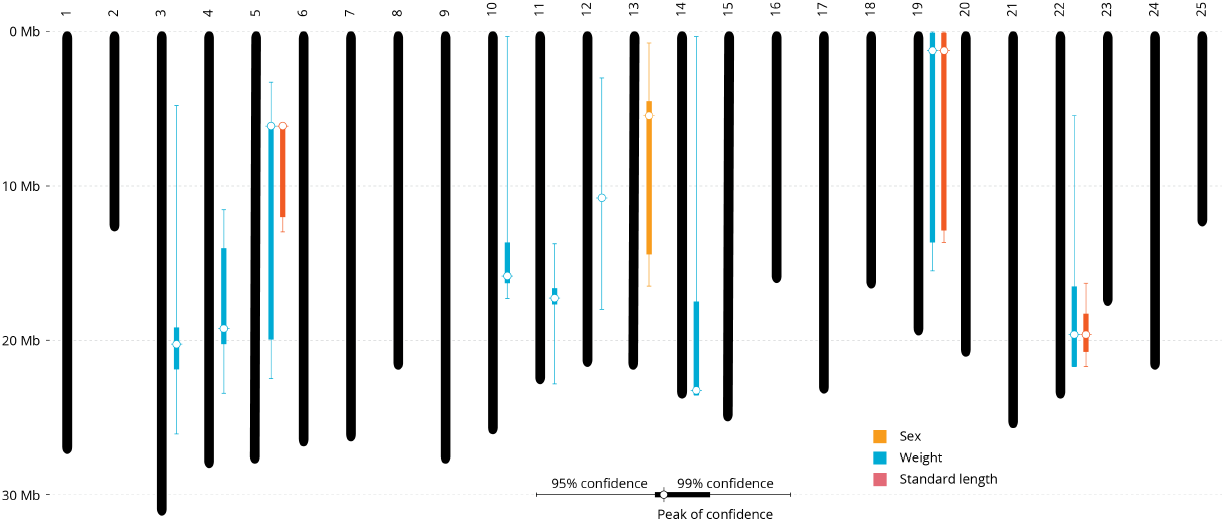
QTL map. Chromosomal locations of highlighted genomic regions for QTLs in this study, including Standard length, fish weight, phenotypic sex and condition factor. The peak locations are located with white circles.

### Prediction and Validation

The combined prediction power of these 177 unique SNP markers (from the sex and weight QTL) was tested by building machine-learning algorithms and using an instance-based k-nearest neighbours’ algorithm (Aha et al., 1991) based on the additive effect of the genotypes at each locus considered. Phenotype prediction power from using these 177 SNPs provided a 99% accuracy for the selection of a desired phenotype (Table 4). The SNP markers defining the QTLs for Weight and Standard length were further investigated to provide a small subset of marker fit for a quick SNP assay. This approach produced a robust combined subset of 29 SNP markers (10 associated to Sex, 14 to Weight and 18 to Standard Length, with weight and standard-length markers overlapping; Supplementary Table S4). When applied to all individuals, the combined prediction power remains over 90% (Table 4).

**Table 4.** Details of the Phenotypic Variation Explained and prediction accuracy for the full SNP dataset and reduced subset. For each trait tested, the subset of SNPs is reported between brackets. The marker subsets overlap. Sex is a binary character, where correct prediction is provided, Weight and Std length are continuous variables where Precision (Correlation) is specified. Subset list is provided in Supplementary Table S4 (29 unique SNP markers).

|  | Marker | Correct prediction / Precision | Mean absolute error |
| --- | --- | --- | --- |
| Sex (All) | 177 | 99.7% | 0.008 |
| Sex (subset) | 10 | 95.5% | 0.0742 |
| Weight (All) | 177 | 99.9% | 10.257 g |
| Weight (subset) | 14 | 92.5% | 67.468 g |
| Std length (All) | 177 | 99.8% | 1.433 mm |
| Std length (subset) | 18 | 91.4% | 16.410 mm |

### Low Density SNP Panel

A total of 177 SNP markers of interest (from QTL regions) and 399 top F_*st*_ SNP markers were combined in a low-density panel to test its usefulness in quantifying and maintaining genetic diversity within the tested population, along with potential use for selection purposes in the future. This final panel of 576 markers successfully delivering informative genotypes was selected (Supplementary Data S2) and evaluated on the previously mentioned 477 samples, showing its usefulness to determine population and family structure, as well as to provide genotypes that can be used for selection purposes (Figure 4).

**Figure 4.**
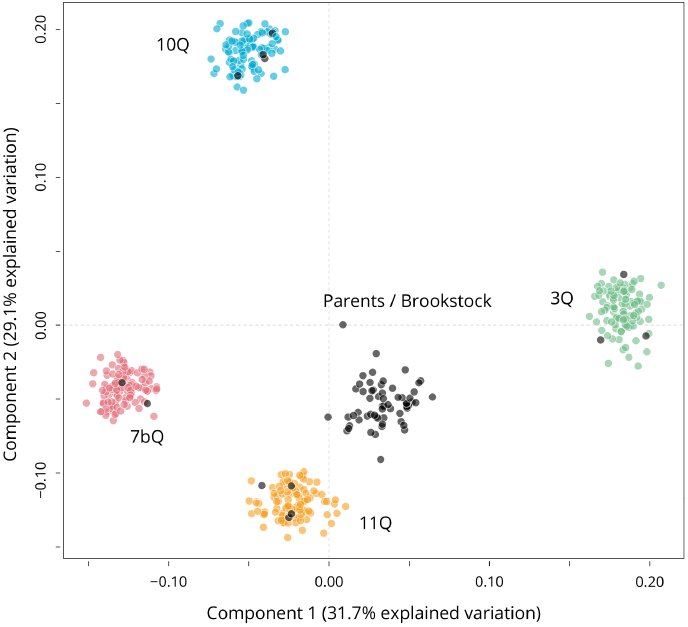
Multidimensional Scaling Analysis results of the validation panel (477 samples) dataset. The first and second components explain 31.7%, and 29.1% of the variation found. Based on 576 SNP markers.

## DISCUSSION

### Importance of generating genomic resources for emerging species

The development of genomic resources for important and emerging aquaculture species is crucial for understanding their biology and paves the way for successful breeding schemes and improved selection (Houston et al., 2020; Robledo et al., 2018). Genomic tools can be particularly advantageous for emerging species such as lumpfish, as it can expedite and improve the accuracy of the selection process for important traits, in addition to establishing breeding programmes.

In this study, genomic markers were developed for a lumpfish stock, with 7,193 informative SNPs being identified following a thorough QC filtering process. This is a significant achievement and represents a valuable starting point for future genomic research on the species. Genomic resources, such as DNA markers, have become an essential component of successful aquaculture production. As a result, many relevant aquaculture species have been targeted for SNP marker development via genotyping-by-sequencing (GBS) or whole genome sequencing (Robledo et al., 2018; Yue and Wang, 2017).

The development of SNP markers has been extensively researched in Atlantic salmon over the last two decades, starting with the screening of a few hundred SNPs, to reach the development and application of numerous SNP arrays containing up to 900K markers (Moen et al., 2008; Gutierrez et al., 2012; Houston et al., 2014; Yáñez et al., 2016; Sinclair-Waters et al., 2020). Sea bream and sea bass are additional examples of species with successful genomic marker development, transitioning from GBS-based SNP identification to the development of medium-density SNP arrays (Palaiokostas et al., 2015, 2016; Peñaloza et al., 2021). This progress exemplifies how commercial interest and production needs can positively stimulate research advancement in aquaculture species, which could serve as a model for lumpfish research given the current demand for cleaner fish.

### GWAS on growth and sex in other species

The increased accessibility to sequencing technologies has made GWAS (and QTL) analyses involving thousands of markers a norm for studying aquaculture and livestock species. This has enabled the identification of significant associations between genomic markers and particular phenotypes, such as growth, sex, disease resistance, and colour, among others, which is a fundamental step towards the selective improvement of stocks. Aquaculture species have been extensively researched for the identification of QTL regions and markers associated with important traits (Yáñez et al., 2023).The present study aimed to identify genomic regions associated with analysed traits, and was successful in this regard (Table 3). The analysis of sex showed the strongest association, with a single major QTL located on chromosome 13 (Figure 2a and Figure 3) being identified. This result is in line with what was recently reported for another lumpfish stock, where chromosome 13 was also identified as the sex chromosome, and the *Amh* gene was suggested as the sex determining gene (Holborn et al., 2022), although the QTL peak position does not exactly match the position of the *Amh* gene in the lumpfish genome. This could be due to many reasons, first the previous study utilized a 70K SNP array for lumpfish, while our analyses were based on 7,193 SNPs, and therefore, even though our results agree with the chromosome location, the lower marker density did not provide enough resolution to identify the specific location of the candidate gene. Nevertheless, a set of 10 SNPs located within this region accurately predicted sex in all samples, giving evidence that the sex determining locus is shared between populations.

Growth rate is a significant trait for improvement in newly domesticated species, and it has been extensively studied in most aquaculture species (Yáñez et al., 2023). Analysis of weight and length in lumpfish showed polygenic involvement, identifying significant associations across many chromosomes, and showing evidence of overlap in QTL regions in chromosomes 5, 19, 14, and 22 (Figure 2b-c, Table 3, and Figure 3). In contrast, the analysis of condition factor (K) did not identify significant associations, most likely due to the round morphology of lumpfish, which makes this index unreliable / uniformative for this species (Garcia de Leaniz et al., 2022). The polygenic nature of growth traits is not surprising, as most aquaculture species show this pattern. Contrary to most reported aquaculture goals, where growth rate QTLs have been largely exploited to increase growth rate, selection for slower growth rate and longer deployment time may be feasible for lumpfish. Grazing efficacy has been negatively correlated with the size of lumpfish and linked to parental/family effects, suggesting that the genetic component can play a significant role in improving growth and grazing (Imsland et al., 2021). The markers identified in this study show promise for the selection of slow-growing fish using a low number of markers (Table 4), and therefore, they have the potential to improve the grazing efficiency of selected stocks.

### Usefulness of findings. MAS and genomic selection applications

The QTL markers identified in this study have great potential to significantly enhance the analysed traits, particularly growth, which has shown average genetic gains of over 10% per generation in some aquaculture species (Gjedrem et al., 2012). Selection to obtain the opposite outcome (slower growth) should be possible at similar rates, particularly with the introduction of genomic resources into the selection process. Our analysis of 177 markers associated with the traits revealed 99% accuracy in predicting the animal’s phenotype, and a selection of only 29 SNPs achieved similar accuracy, thus opening the possibility of using low-density SNP panels, such as the one described in this study, to provide practical genomic resources at a lower cost without sacrificing selection power.

The results of this study demonstrate that a panel of 576 markers can determine family structure and accurately predict slow growth phenotypes, and sex-associated markers can accurately distinguish the sex of individuals, which is particularly beneficial for selecting broodstock at early stages (Table 4 and Figure 4). Furthermore, these genomic resources can be used to determine relatedness, population structure, genetic variation, and inform genomic selection (Kriaridou et al., 2020). While further analyses are necessary to explore the SNP panel’s ability to differentiate the geographical origin of lumpfish populations and test the application of genomic selection for improving selection schemes, the findings provide significant insight into the genetic control of important traits in lumpfish.

Overall, the developed genomic resources offer great potential for facilitating the development of breeding programs for lumpfish and selection based on genomic information. Our study sheds light on the genetic factors influencing growth and sex in lumpfish and highlights the potential of low-density SNP panels as a cost-effective and powerful tool for genomic selection in aquaculture.

## AUTHOR CONTRIBUTIONS

Conceptualisation: A.D., I.L., H.M., A.P.G., P.K., S.-L.C.S. and M.B.; Data curation: P.K. and M.B.; Formal analysis: A.P.G., P.K. and M.B.; Funding acquisition: A.D., H.M., S.-L.C.S., I.L. and T.C.R.; Investigation: A.P.G., M.B. and P.K.; Methodology: A.D., A.P.G., J.B.T., M.B., P.K. and S.-L.C.S.; Project administration: A.P.G.; Resources: A.D., A.P.G., S.-L.C.S. and T.C.R.; Supervision: A.P.G. and M.B.; Validation: M.B. and P.K.; Writing - original draft: A.P.G. and M.B.; Writing - review & editing: A.D., A.P.G., E.J.F., I.L., H.M., J.B.T., M.B., P.K., S.-L.C.S. and T.C.R.

## INSTITUTIONAL REVIEW

Ethical approval for the study was granted in accordance with the Norwegian state Mattilsynet program and in accordance with the UK Animals (Scientific Procedures) Act 1986 Amended Regulations (SI 2012/3039), as well as approved by the Animal Welfare and Ethical Review Body of the University of Stirling (AWERB (18 19) 032).

## DATA AVAILABILITY

All the raw reads sequence data obtained in this study were deposited at the European Bioinformatics Institute (EBI) Sequence Read Archive (SRA) project ID PRJEB38408

## FUNDING

This research was funded by EU funded AquaExcel 2020, Trans-national access grant (AE100022), the Biotechnology and Biological Sciences Research Council and Natural Environment Research Council [grant number BB/S004416/1] and the Sustainable Aquaculture Innovation Centre.

## CONFLICTS OF INTEREST

T.C. and E.J.F. are employed by Otter Ferry SeaFish Ltd, a funding partner of the research project. The other authors declare that the research was conducted in the absence of any commercial or financial relationships that could be construed as a potential conflict of interest.

## SUPPLEMENTARY MATERIAL

**Table S1**. Details of the samples used, metadata, barcodes and read numbers.

**Table S2**. Details of the samples used for the panel validation.

**Table S3**. Details of the markers, genomic location, *p*-value of association with Standard length, Weight, and Sex.

**Table S4**. Details of the subset of SNP markers.

**Data S1**. Genotypes of the 536 samples and 7,193 markers. Each marker is located on the GCA 009769545.1 assembly (VCF).

**Data S2**. Details of the 576 panel SNP markers, genomic location (BED).

## Notes

### Competing Interest Statement

The authors have declared no competing interest.

## REFERENCES

Aha, D. W., Kibler, D., and Albert, M. K. (1991). Instance-based learning algorithms. Mach. Learn., 6(1):37–66.

Bolton-Warberg, M. (2018). An overview of cleaner fish use in ireland. J. Fish Dis., 41(6):935–939.

Broman, K. W., Gatti, D. M., Simecek, P., Furlotte, N. A., Prins, P., Sen, Ś., Yandell, B. S., and Churchill, G. A. (2018). R/qtl2: Software for mapping quantitative trait loci with highdimensional data and multiparent populations. Genetics, 211(2):495–502.

Broman, K. W. and Sen, S. (2009). A guide to QTL mapping with R/qtl. Statistics for biology and health. Springer, New York, NY.

Brooker, A. J., Papadopoulou, A., Gutierrez, C., Rey, S., Davie, A., and Migaud, H. (2018). Sustainable production and use of cleaner fish for the biological control of sea lice: recent advances and current challenges. Vet. Rec., 183(12):383–383.

Brown, J. K., Taggart, J. B., Bekaert, M., Wehner, S., Palaiokostas, C., Setiawan, A. N., Symonds, J. E., and Penman, D. J. (2016). Mapping the sex determination locus in the hpuku (Polyprion oxygeneios) using ddRAD sequencing. BMC Genomics, 17:448.

Catchen, J. M., Amores, A., Hohenlohe, P., Cresko, W., and Postlethwait, J. H. (2011). Stacks: Building and genotyping loci de novo from short-read sequences. G3, 1(3):171–182.

Costello, M. J. (2006). Ecology of sea lice parasitic on farmed and wild fish. Trends Parasitol., 22(10):475–483.

Costello, M. J. (2009). The global economic cost of sea lice to the salmonid farming industry. J. Fish Dis., 32(1):115–118.

Davenport, J. (1985). Synopsis of biological data on the lumpsucker, Cyclopterus lumpus (Linnaeus, 1758). xumber no. 147 in FAO fisheries synopsis. Food and Agriculture Organization of the United Nations, Rome, Italy.

Garcia de Leaniz, C., Gutierrez Rabadan, C., Barrento, S. I., Stringwell, R., Howes, P. N., Whittaker, B. A., Minett, J. F., Smith, R. G., Pooley, C. L., Overland, B. J., Biddiscombe, L., Lloyd, R., Consuegra, S., Maddocks, J. K., Deacon, P. T. J., Jennings, B. T., Rey Planellas, S., Deakin, A., Moore, A. I., Phillips, D., Bardera, G., Castanheira, M. F., Scolamacchia, M., Clarke, N., Parker, O., Avizienius, J., Johnstone, M., and Pavlidis, M. (2022). Addressing the welfare needs of farmed lumpfish: Knowledge gaps, challenges and solutions. Rev. Aquac., 14(1):139–155.

Gjedrem, T., Robinson, N., and Rye, M. (2012). The importance of selective breeding in aquaculture to meet future demands for animal protein: A review. Aquaculture, 350-353:117–129.

González, J. R., Armengol, L., Solé, X., Guinó, E., Mercader, J. M., Estivill, X., and Moreno, V. (2007). SNPassoc: an R package to perform whole genome association studies. Bioinformatics, 23(5):644–645.

Gruber, B., Unmack, P. J., Berry, O. F., and Georges, A. (2018). dartR: An R package to facilitate analysis of SNP data generated from reduced representation genome sequencing. Mol. Ecol. Resour., 18(3):691–699.

Gutierrez, A. P., Lubieniecki, K. P., Davidson, E. A., Lien, S., Kent, M. P., Fukui, S., Withler, R. E., Swift, B., and Davidson, W. S. (2012). Genetic mapping of quantitative trait loci (QTL) for body-weight in Atlantic salmon (Salmo salar) using a 6.5 K SNP array. Aquaculture, 358-359:61–70.

Holborn, M. K., Einfeldt, A. L., Kess, T., Duffy, S. J., Messmer, A. M., Langille, B. L., Brachmann, M. K., Gauthier, J., Bentzen, P., Knutsen, T. M., Kent, M., Boyce, D., and Bradbury, I. R. (2022). Reference genome of lumpfish Cyclopterus lumpus linnaeus provides evidence of male heterogametic sex determination through the AMH pathway. Mol. Ecol. Resour., 22(4):1427–1439.

Houston, R. D., Bean, T. P., Macqueen, D. J., Gundappa, M. K., Jin, Y. H., Jenkins, T. L., Selly, S. L. C., Martin, S. A. M., Stevens, J. R., Santos, E. M., Davie, A., and Robledo, D. (2020). Harnessing genomics to fast-track genetic improvement in aquaculture. Nat. Rev. Genet., 21(7):389–409.

Houston, R. D., Taggart, J. B., Cézard, T., Bekaert, M., Lowe, N. R., Downing, A., Talbot, R., Bishop, S. C., Archibald, A. L., Bron, J. E., Penman, D. J., Davassi, A., Brew, F., Tinch, A. E., Gharbi, K., and Hamilton, A. (2014). Development and validation of a high density SNP genotyping array for Atlantic salmon (Salmo salar). BMC Genomics, 15:90.

Imsland, A. K., Reynolds, P., Nytrø, A. V., Eliassen, G., Hangstad, T. A., Jónsdóttir, Ó. D., Emaus, P.-A., Elvegård, T. A., Lemmens, S. C., Rydland, R., and Jonassen, T. M. (2016). Effects of lumpfish size on foraging behaviour and co-existence with sea lice infected atlantic salmon in sea cages. Aquaculture, 465:19–27.

Imsland, A. K. D., Hanssen, A., Nytrø, A. V., Reynolds, P., Jonassen, T. M., Hangstad, T. A., Elvegård, T. A., Urskog, T. C., and Mikalsen, B. (2018). It works! lumpfish can significantly lower sea lice infestation in large-scale salmon farming. Biol. Open, 7(9):bio036301.

Imsland, A. K. D., Reynolds, P., Hangstad, T. A., Kapari, L., Maduna, S. N., Hagen, S. B., Jónsdóttir, Ó. D. B., Spetland, F., and Lindberg, K. S. (2021). Quantification of grazing efficacy, growth and health score of different lumpfish (Cyclopterus lumpus L.) families: Possible size and gender effects. Aquaculture, 530:735925.

Jolliffe, I. T. and Cadima, J. (2016). Principal component analysis: a review and recent developments. Philos Trans A Math Phys Eng Sci, 374(2065):20150202.

Kriaridou, C., Tsairidou, S., Houston, R. D., and Robledo, D. (2020). Genomic prediction using low density marker panels in aquaculture: Performance across species, traits, and genotyping platforms. Front. Genet., 11:124.

Langmead, B. and Salzberg, S. L. (2012). Fast gapped-read alignment with Bowtie 2. Nat. Methods, 9(4):357–359.

Moen, T., Hayes, B., Baranski, M., Berg, P. R., Kjøglum, S., Koop, B. F., Davidson, W. S., Omholt, S. W., and Lien, S. (2008). A linkage map of the atlantic salmon (Salmo salar) based on EST-derived SNP markers. BMC Genomics, 9:223.

Nash, R. D. M., Valencia, A. H., and Geffen, A. J. (2006). The origin of fulton’s condition factor - setting the record straight. Fisheries, 31(5):236–238.

Norwegian Directorate of Fisheries (2022). Sale of farmed cleaner fish 2012–2021.

Overton, K., Barrett, L. T., Oppedal, F., Kristiansen, T. S., and Dempster, T. (2020). Sea lice removal by cleaner fish in salmon aquaculture: a review of the evidence base. Aquacult. Environ. Interact., 12:31–44.

Palaiokostas, C., Bekaert, M., Taggart, J. B., Gharbi, K., McAndrew, B. J., Chatain, B., Penman, D. J., and Vandeputte, M. (2015). A new SNP-based vision of the genetics of sex determination in European sea bass (Dicentrarchus labrax). Genet. Sel. Evol., 47(1):68.

Palaiokostas, C., Ferraresso, S., Franch, R., Houston, R. D., and Bargelloni, L. (2016). Genomic prediction of resistance to Pasteurellosis in gilthead sea bream (Sparus aurata) using 2b-RAD sequencing. G3, 6(11):3693–3700.

Peñaloza, C., Manousaki, T., Franch, R., Tsakogiannis, A., Sonesson, A., Aslam, M., Allal, F., Bargelloni, L., Houston, R., and Tsigenopoulos, C. (2021). Development and testing of a combined species SNP array for the European seabass (Dicentrarchus labrax) and gilthead seabream (Sparus aurata). Genomics, 113(4):2096–2107.

Powell, A., Treasurer, J. W., Pooley, C. L., Keay, A. J., Lloyd, R., Imsland, A. K., and Garcia de Leaniz, C. (2018). Use of lumpfish for sea-lice control in salmon farming: challenges and opportunities. Rev. Aquac., 10(3):683–702.

Purcell, S., Neale, B., Todd-Brown, K., Thomas, L., Ferreira, M. A., Bender, D., Maller, J., Sklar, P., de Bakker, P. I., Daly, M. J., and Sham, P. C. (2007). PLINK: a tool set for whole-genome association and population-based linkage analyses. Am. J. Hum. Genet., 81(3):559–75.

R Core Team (2022). R: A language and environment for statistical computing. https://www.r-project.org.

Regan, T., Bean, T. P., Ellis, T., Davie, A., Carboni, S., Migaud, H., and Houston, R. D. (2021). Genetic improvement technologies to support the sustainable growth of UK aquaculture. Rev. Aquac., 13(4):1958–1985.

Robledo, D., Palaiokostas, C., Bargelloni, L., Martínez, P., and Houston, R. (2018). Applications of genotyping by sequencing in aquaculture breeding and genetics. Rev. Aquac., 10(3):670–682.

Sinclair-Waters, M., Ødegård, J., Korsvoll, S. A., Moen, T., Lien, S., Primmer, C. R., and Barson, N. J. (2020). Beyond large-effect loci: large-scale GWAS reveals a mixed large-effect and polygenic architecture for age at maturity of Atlantic salmon. Genet. Sel. Evol., 52:9.

Witten, I. H., Frank, E., Hall, M. A., and Pal, C. J. (2017). The WEKA workbench, pages 553–571. Morgan Kaufmann, Cambridge, MA, fourth edition.

Yáñez, J. M., Barría, A., López, M. E., Moen, T., Garcia, B. F., Yoshida, G. M., and Xu, P. (2023). Genome-wide association and genomic selection in aquaculture. Rev. Aquac., 15(2):645–675.

Yáñez, J. M., Naswa, S., López, M. E., Bassini, L., Correa, K., Gilbey, J., Bernatchez, L., Norris, A., Neira, R., Lhorente, J. P., Schnable, P. S., Newman, S., Mileham, A., Deeb, N., Genova, A. D., and Maass, A. (2016). Genomewide single nucleotide polymorphism discovery in Atlantic salmon (Salmo salar): validation in wild and farmed American and European populations. Mol. Ecol. Resour., 16(4):1002–1011.

Yue, G. and Wang, L. (2017). Current status of genome sequencing and its applications in aquaculture. Aquaculture, 468(1):337–347.

Zheng, X., Levine, D., Shen, J., Gogarten, S. M., Laurie, C., and Weir, B. S. (2012). A highperformance computing toolset for relatedness and principal component analysis of SNP data. Bioinformatics, 28(24):3326–3328.

